# Theoretical Considerations and Empirical Predictions of the Pharmaco- and Population Dynamics of Heteroresistance

**DOI:** 10.1101/2023.09.21.558832

**Authors:** Bruce R. Levin, Brandon A. Berryhill, Teresa Gil-Gil, Joshua A. Manuel, Andrew P. Smith, Jacob E. Choby, Dan I. Andersson, David S. Weiss, Fernando Baquero

**Affiliations:** Department of Biology, Emory University; Atlanta, Georgia, 30322, USA; Emory Antibiotic Resistance Center; Atlanta, Georgia, 30322, USA; Program in Microbiology and Molecular Genetics, Graduate Division of Biological and Biomedical Sciences, Laney Graduate School, Emory University; Atlanta, GA, 30322, USA; Emory Vaccine Center; Atlanta, Georgia, 30322, USA; Department of Medical Biochemistry and Microbiology, Uppsala University, Uppsala, SE-75123, Sweden; Division of Infectious Diseases, Department of Medicine, Emory University School of Medicine; Atlanta, GA, 30322, USA; Georgia Emerging Infections Program, Georgia Department of Public Health; Atlanta, GA, 30322, USA; Servicio de Microbiología, Hospital Universitario Ramón y Cajal, Instituto Ramón y Cajal de Investigación Sanitaria, and Centro de Investigación Médica en Red, Epidemiología y Salud Pública (CIBERESP) Madrid, Spain

**Keywords:** Microbiology, Heteroresistance, Antibiotic Resistance, Pharmacodynamics, Mathematical Modeling

## Abstract

Antibiotics are considered one of the most important contributions to clinical medicine in the last 100 years. Due to the use and overuse of these drugs, there have been increasing frequencies of infections with resistant pathogens. One form of resistance, heteroresistance, is particularly problematic; pathogens appear sensitive to a drug by common susceptibility tests. However, upon exposure to the antibiotic, resistance rapidly ascends, and treatment fails. To quantitatively explore the processes contributing to the emergence and ascent of resistance during treatment and the waning of resistance following cessation of treatment, we develop two distinct mathematical and computer-simulations models of heteroresistance. In our analysis of the properties of these models, we consider the factors that determine the response to antibiotic-mediated selection. In one model, heteroresistance is progressive, with each resistant state sequentially generating a higher resistance level. In the other model, heteroresistance is non-progressive, with a susceptible population directly generating populations with different resistance levels. The conditions where resistance will ascend in the progressive model are narrower than those of the non-progressive model. The rates of reversion from the resistant to the sensitive states are critically dependent on the transition rates and the fitness cost of resistance. Our results demonstrate that the standard test used to identify heteroresistance is insufficient. The predictions of our models are consistent with empirical results. Our results demand a reevaluation of the definition and criteria employed to identify heteroresistance. We recommend the definition of heteroresistance should include a consideration of the rate of return to susceptibility.

**Significance Statement:** This mathematical modeling and computer-simulation study quantitatively explores two broadly different, previously undescribed, classes of heteroresistance. In our analysis of the properties of these models, we consider the response of heteroresistant populations to antibiotic exposure, focusing on the conditions where heteroresistance could lead to clinical treatment failure. We also provide novel consideration to the rate of reversion from a resistant to sensitive state. Our analysis illustrates the need to include the reversion rate from resistant to sensitive in the definition of heteroresistance and questions the sufficiency of the method currently used to identify heteroresistance.

## Introduction

Pathogens resistant to existing antibiotics are a significant and increasing source of morbidity and mortality for humans and domestic animals (1, 2). Fundamental to the effective treatment of bacterial infections is choosing an antibiotic to which the pathogen is susceptible. The level of susceptibility is readily estimated by culture methods, both through automation via BioMerieux’s VITEK and similar devices (3-6), as well as by non-automated methods such as disk diffusion and Epsilon-diffusion tests (7, 8). By these methods, bacteria are classified as susceptible, intermediate, or resistant according to the international consensus guidelines from the Clinical and Laboratory Standards Institute (CLSI) and the European Committee on Antimicrobial Susceptibility Testing (EUCAST). These categorical descriptions determine whether an antibiotic will or will not be used for treatment. If an isolate appears susceptible to an antibiotic by these criteria, the drug would be presumed to be effective in treating infections with that pathogen. These *in vitro* susceptibility estimates are not sufficient as measures of antibiotic susceptibility if the treated bacteria are heteroresistant to that drug.

A population of bacteria which is heteroresistant often appears susceptible to an antibiotic as assessed by the standard methods described above, but quickly becomes resistant upon confrontation with that drug due to the selection for and ascent of minority resistant populations. Heteroresistance (HR) is typically defined by the presence of one or more sub-populations at a frequency greater than 10^-7^ with a resistance level that crosses the breakpoint at or greater than 8 times the susceptible main population (9). The canonical test for the presence of these sub-populations, and thus for HR, is a Population Analysis Profile (PAP) test (10, 11). This protocol tests for bacterial growth at different concentrations of an antibiotic, thus revealing the presence or absence of resistant sub-populations.

HR is clinically and epidemiology problematic due to the inherent instability of resistance. Within short order of the removal of the antibiotic, heteroresistant populations once again appear susceptible to the treating antibiotic by conventional testing procedures. This effect is most profound when considering the transmission of heteroresistant populations between individuals. Patients with heteroresistant infections transmit these seemingly antibiotic-susceptible bacteria to other patients, who may then fail treatment with the drug for which the bacteria are HR. This instability of resistance is intrinsic to HR but is not currently part of the definition and thus is considered in few reports of HR (12).

In this report, we develop and analyze the properties of two mathematical and computer-simulation models that represent two extreme cases of HR, which we call progressive and non-progressive. Using these models, we explore the pharmaco- and population dynamic processes responsible for HR and the factors which contribute to the instability of HR. The parameters of these models can be estimated, and the hypotheses generated therefrom tested and rejected *in vitro* and *in vivo*.

## Results

### Models of Heteroresistance

We open this consideration of the pharmaco- and population dynamics of HR with a description of the two mathematical models employed. For both models of HR, we assume a Hill function for the relationship between the concentration of the antibiotic, the concentration of the limiting resource, and the rates of growth and death of the bacteria, known as the pharmacodynamics (13-15).

#### Pharmacodynamics

In accord with the Hill function, the rate of growth or death of bacteria exposed to a given antibiotic concentration is given by Equation 1.

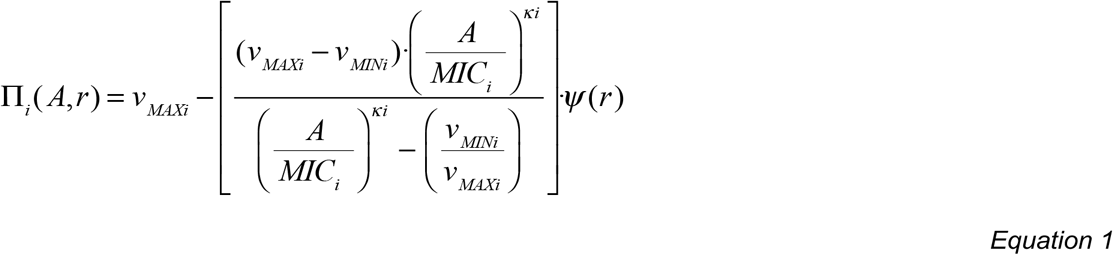

Where *A* in µg/mL is the antibiotic concentration and *r* in µg/mL is the concentration of the resource which limits the growth of the population. *v*_*MAXi*_ is the maximum growth rate in cells per hour of the bacteria of state *i*, where *v*_*MAXi*_*>0. v*_*MINi*_ is the minimum growth rate per cell per hour, which is the maximum death rate when exposed to the antibiotic, where *v*_*MINi*_<0. *MIC*_*i*_ is the minimum inhibitory concentration of the antibiotic for the bacteria of state *i* in µg/mL. *κi* is the Hill coefficient for bacteria of state *i*. The greater the value of *κi*, the more acute the function. The function, 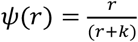, is the rate of growth in the absence of the antibiotic, where *k* is the resource concentration in µg/mL when the growth rate is half of its maximum value. ψ(r) measures the physiological state of the bacteria; as the resource concentration declines the cells grow slower. We show in Supplemental Figure 1 the Hill functions for four different bacterial populations with varying MICs and maximum growth rates.

#### Diagrams of the Heteroresistance Models

The two models of heteroresistance used here are depicted in Figure 1. In the progressive model (Figure 1A) the increasingly resistant states are generated by a transition from a less resistant state to a more resistant state, and the more resistant states generate the less resistant states sequentially. In the non-progressive model (Figure 1B) the different resistant states are generated directly by a transition from the susceptible state, and the more resistant states transition directly back to the most sensitive state.

**Figure 1.**
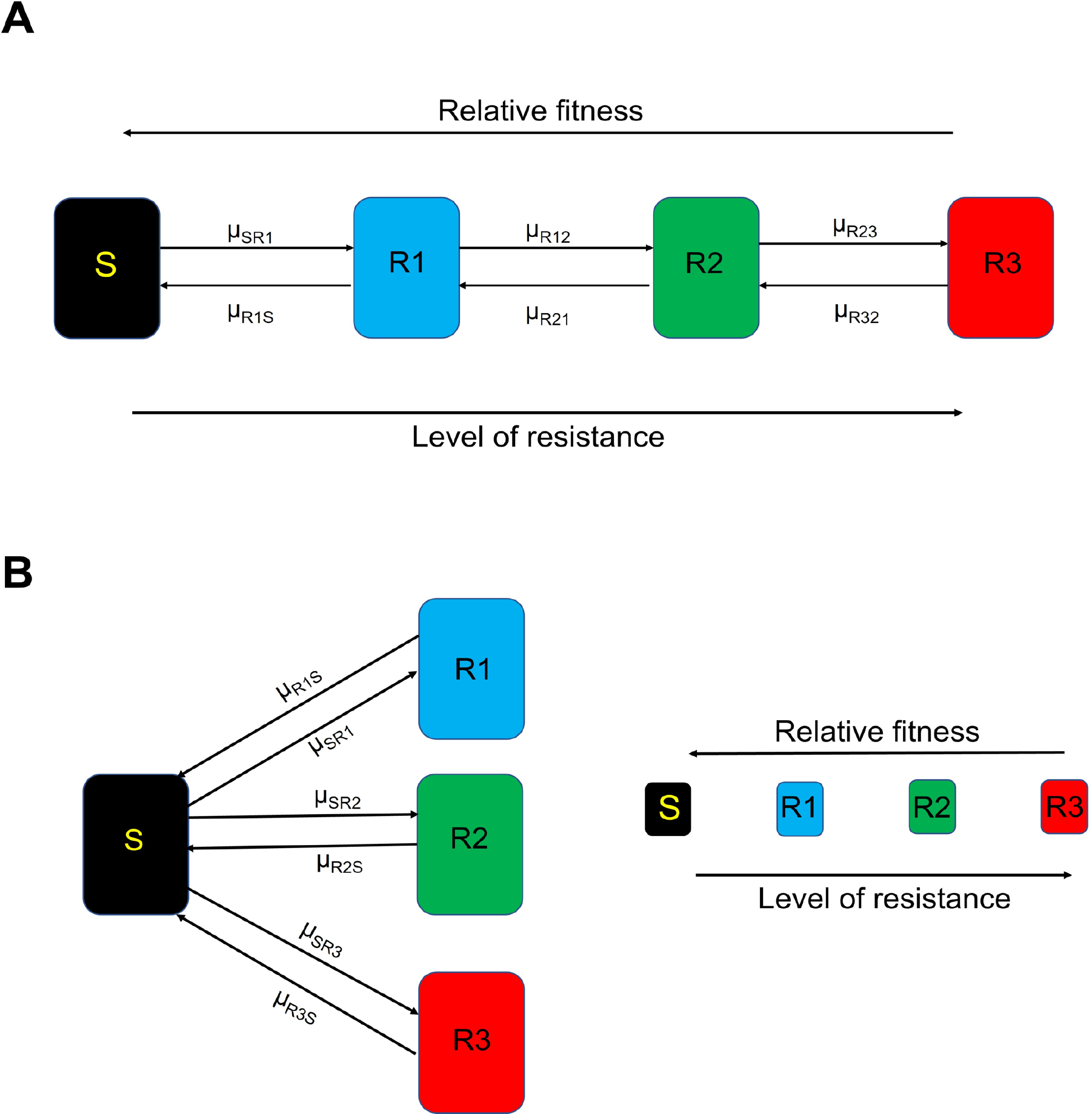
Diagram of the two models of HR. S (black) is the most antibiotic-sensitive state and the state with the highest fitness. We assume the level of antibiotic resistance increases as the fitness decreases from state R1 (blue) to R2 (green) to R3 (red). Transitions occur between states at potentially different rates of µ (where µ_ij_ is the transition from i to j). Panel A is a diagram of the progressive model and panel B is a diagram of the non-progressive model.

#### The Progressive Model

In this model (Figure 1A), the bacteria transition between four different states: sensitive, S, and increasingly resistant, R1, R2, R3, which are the designations and densities in cells per mL of bacteria of these different states. Cells of the S state transition to R1, R1 transitions to R2, and R2 transitions to R3 at rates µ_S1_, µ_R12_, and µ_R23_ per cell per hour, respectively. Cells of resistant states progressively transition to the less resistant states, R3 to R2, R2 to R1, and R1 to S, with rates µ_R32_, µ_R21_, and µ_R1S_ per cell per hour. We simulate these transitions with a Monte Carlo process (16). A random number x (0 ≤ x ≤1) from a rectangular distribution is generated. If x is less than the product of the number of cells in the generating state (S, the density time the volume of the vessel, Vol), the transition rate (µ) and the step size (dt) of the Euler method (17) employed for solving differential equation, for example if x < S*µ_SR1_*dt*Vol, then MSR1 cells are added to the R1 population and removed from the S population where MSR1=1/(dt*Vol). With these definitions, assumptions, and the parameters defined and presented in Supplemental Table 1, the rates of change in the densities of the different populations will be given by:

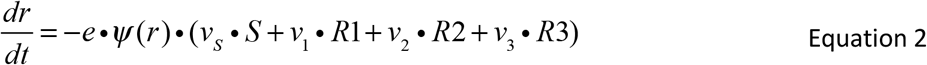

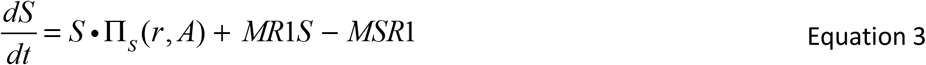

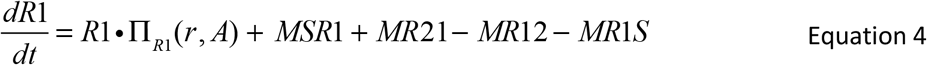

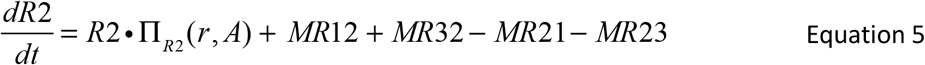

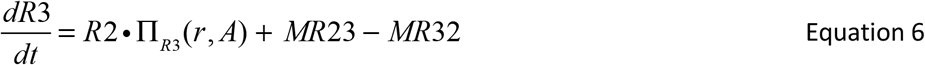

#### The Non-progressive Model

In this model (Figure 1B) all the resistant states, R1, R2, and R3 are derived from the susceptible state and, by transition, return directly to the susceptible state, S. The rates of transition from state S are respectively µ_SR1_, µ_SR2_, and µ_SR3_ per cell per hour. The rates of return to the susceptible state are µ_R1S_, µ_R2S_, and µ_R3S_ per cell per hour. The transitions between states are via a Monte Carlo process (16), using a routine like that for the progressive model. When transients from S to the different R states are generated (MSR1, MSR2, and MSR3), 1/(dt*Vol) are added to the R1, R2 and R3 populations and are removed from the S population. When transients from the R1, R2, and R3 populations are generated (MR1S, MR2S and MR3S), 1/(dt*Vol) are added to the S population and removed from the R1, R2, and R3 populations, respectively. With these definitions, assumptions, and the parameters defined and presented in Table S1 the rates of change in the densities of the different populations are given by:

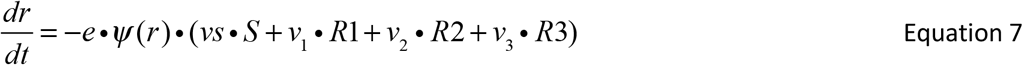

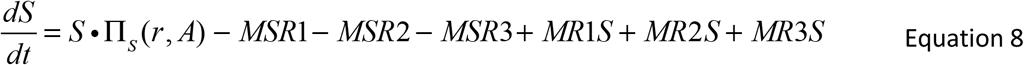

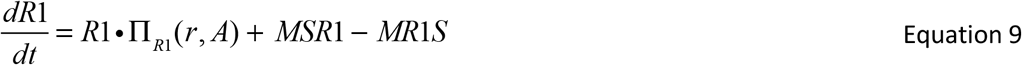

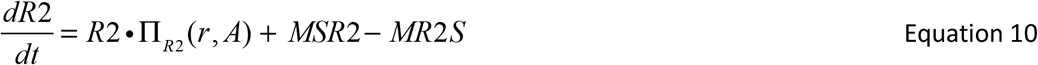

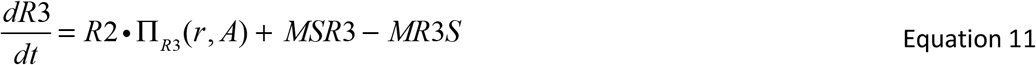

### Simulated Population Dynamics of Heteroresistance

Here we consider the population dynamics of heteroresistance with the distributions of the different resistant states generated from single cells grown up to full densities for the progressive and non-progressive models with four transition rates (Figure 2). We further consider a greater range of transition rates for the non-progressive model to determine the minimum rate for which we generate sufficiently large minority populations in Supplemental Figure 2A-C.

**Figure 2.**
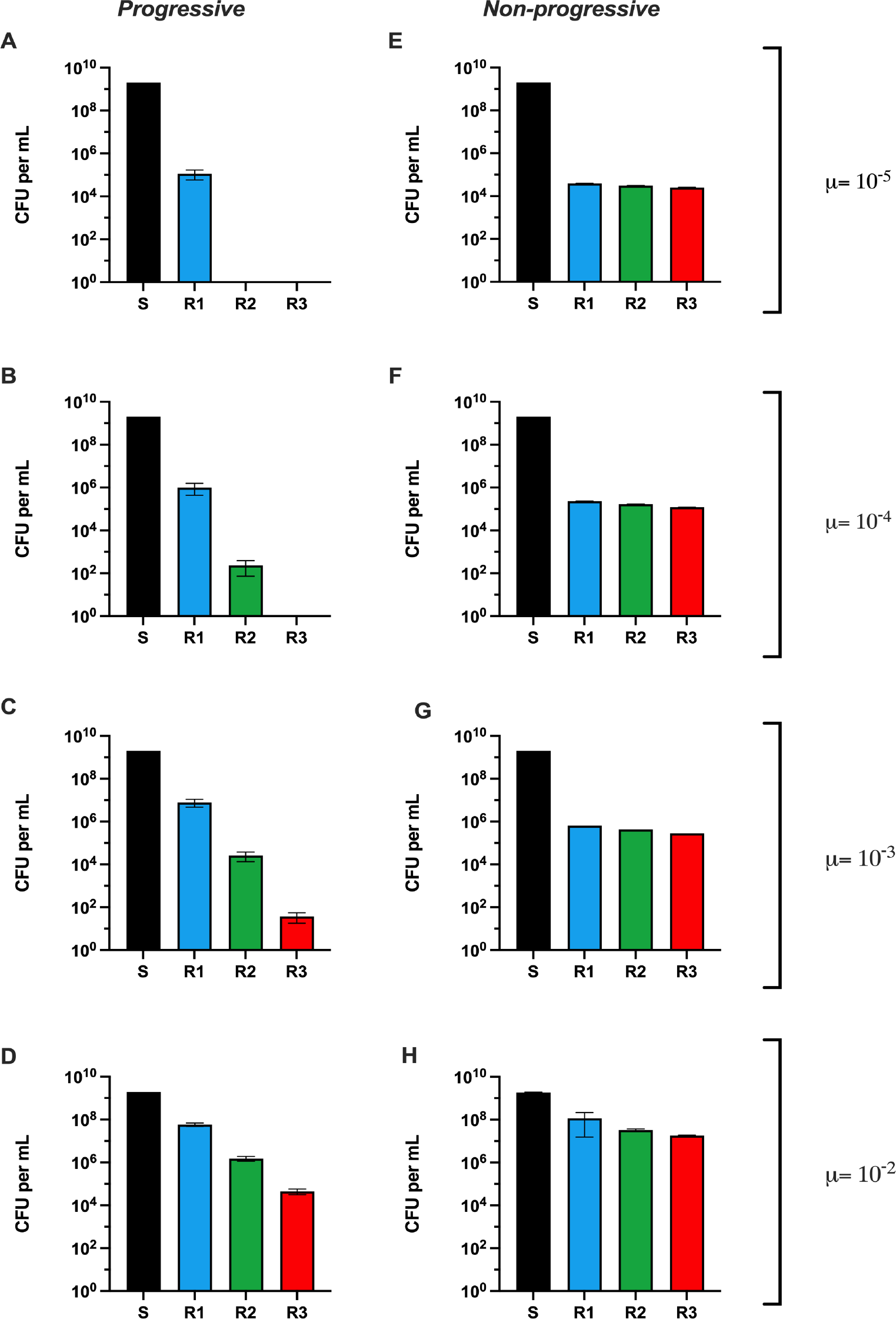
Distribution of stationary phase densities when grown up from a single cell of S. Shown on the left (A, B, C, and D) are the means and standard deviations of the stationary phase densities of the S (black), R1 (blue), R2 (green), and R3 (red) populations from five independent runs with the progressive model with different transition rates, µ=10^-5^, 10^-4^, 10^-3^, and 10^-2^ per cell per hour for A, B, C, and D, respectively. On the right, E, F, G, and H are the corresponding distributions for runs made with the non-progressive model with these respective transition rates.

For the progressive model, only in the runs with the highest transition rates, µ=10^-2^ and µ=10^-3^ per cell per hour, is the subpopulation with the highest resistance level, R3, present. A very different situation obtains for the non-progressive model, as at every transition rate the R3 population is present. We also consider the effect that the relative fitness cost of each state has on these stationary phase densities (Supplemental Figure 3) and find modest differences in these distributions.

Using these same parameters for both models, another difference can be seen between the progressive and non-progressive models in the Population Analysis Profile (PAP) tests of each (Figure 3). For these PAP tests, we calculate the ratio of the number of cells generated at a particular antibiotic concentration compared to the number of cells present when there is no antibiotic (N(A)/N(0)) for 0, 1, 2, 4, 8, and 16 times the MIC of the susceptible population.

**Figure 3.**
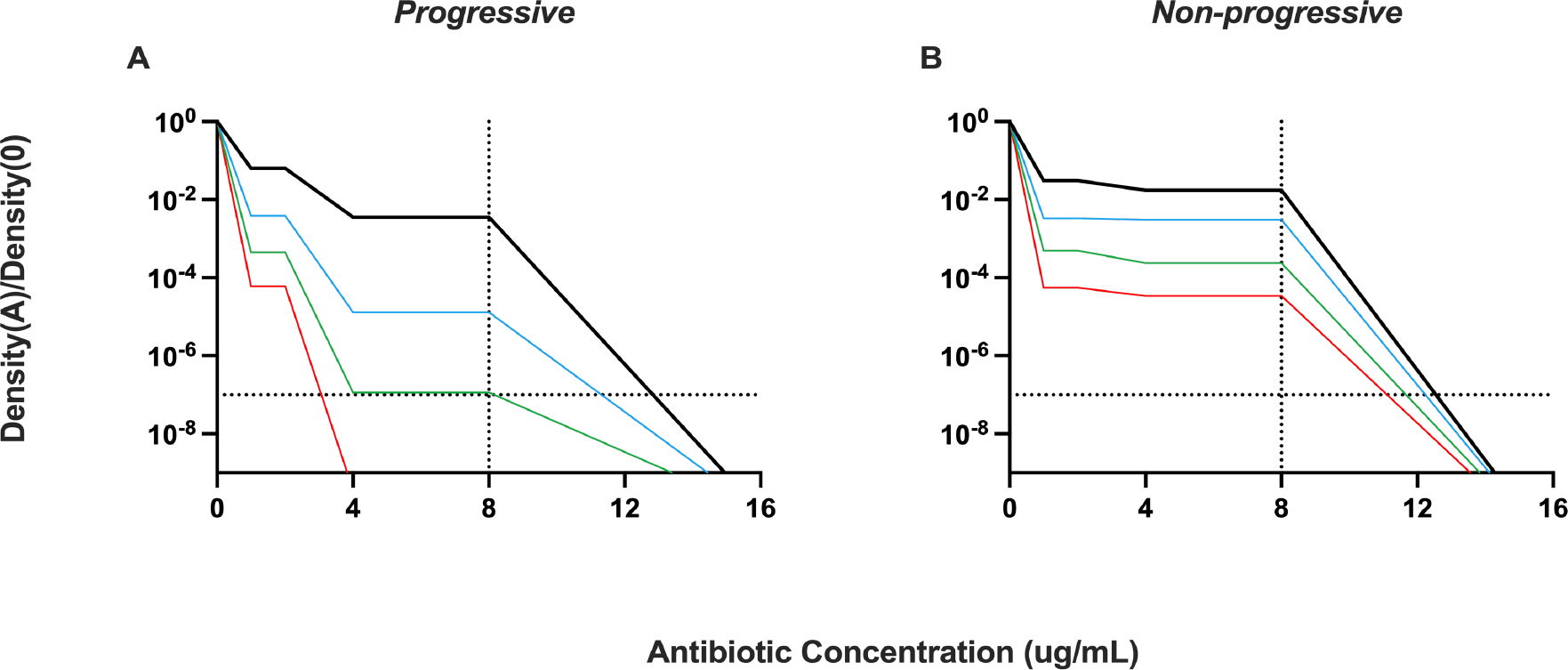
Population Analysis Profile (PAP) tests. The ratio of the density of the number of bacteria surviving at an antibiotic concentration relative to that surviving in the absence of the antibiotic for different transition rates. Black µ=10^-2^, blue µ=10^-3^, green µ=10^-4^, and red µ=10^-5^ per cell per hour. Panel A is the PAP test using the progressive model and Panel B is the PAP test using the non-progressive model.

The PAP test results anticipated from the progressive model are very different than those anticipated from the non-progressive model, two extreme HR cases. The presence of four sub-populations with different MICs is apparent from the PAP test of the progressive model with the parameters considered. For the non-progressive model, the differences in the relative densities of the sub-populations are too low to be detected by a PAP test performed in the lab. In general, the plateaus shown in Figure 3 A and B are sharper and more dramatic than would be seen in the lab. This is a consequence of having only four resistant states. Moreover, using the standard HR criteria of having a sub-population with an MIC of >8 times at a frequency of at least 10^-7^, the progressive model only meets these criteria at high transition rates (exceeding 10^-4^). On the other hand, the non-progressive model meets these criteria at transition rates as low as 10^-7^ (Supplemental Figure 2D).

To explore how these models differ in their response to antibiotic treatment, we follow the changes in the densities and MICs of heteroresistant populations exposed to two antibiotic concentrations (5 µg/mL and 10 µg/mL corresponding to 5x and 10x the MIC of the susceptible population) when µ=10^-2^ and µ=10^-5^ per cell per hour (Supplemental Figure 4). We initiate these simulations with 1/100 of the stationary phase densities of the different states anticipated from the heteroresistant populations depicted in Figure 2D and A and Figure 2H and E for the progressive and non-progressive models of heteroresistance, respectively. In Supplemental Figure 5, we consider the effect that the fitness cost of resistance has on these dynamics and find the effect modest at best, just slowing the response time to the antibiotic.

There are apparent differences in the bacterial response to antibiotics between the progressive and non-progressive models of HR. For both models, when µ=10^-2^ per cell per hour, the R3 population comes to dominate and the MIC increases to the maximum (15 µg/mL), though with the higher concentration of the drug, it takes longer for the R3 population to become dominant. With the lower transition rate of µ=10^-5^ per cell per hour, at 5 µg/mL of antibiotic the R2 population comes to dominate in the progressive model and the R3 population remains minor. At this same transition rate and at 10 µg/mL, resistance does not evolve, and the bacterial populations are lost. In both cases, the MIC does increase but does not go to the maximum value. For these conditions, in the non-progressive model, sub-populations are always able to respond to the antibiotic and are never eliminated.

Upon removal of the antibiotic, the heteroresistant bacterial population reverts to the sensitive state. This reversion is the case for both the progressive and non-progressive models of HR considered here. To illustrate this and elucidate the relative contributions of the rates of transition between states and the fitness cost of resistance (as measured by the growth rates) to the dynamics and the time needed to restore susceptibility, we use serial transfer forms of the progressive and non-progressive versions of the HR models. In these simulations, the populations are grown for 24 hours, diluted by a factor of 100, and fresh resources added. In Figure 4, we present the results of simulations of the changes in the densities of the susceptible and resistant populations as well as the change in average MIC in serial transfer following the removal of the antibiotics. These serial transfer simulations were initiated with 10^7^ bacteria per mL of the highest resistance level, R3. We consider two major conditions: one where the fitness cost of resistance is high and another where the fitness cost of resistance is low. In the supplemental materials we consider the dynamics of reversion when a set of even higher fitness costs are used (Supplemental Figure 6).

**Figure 4.**
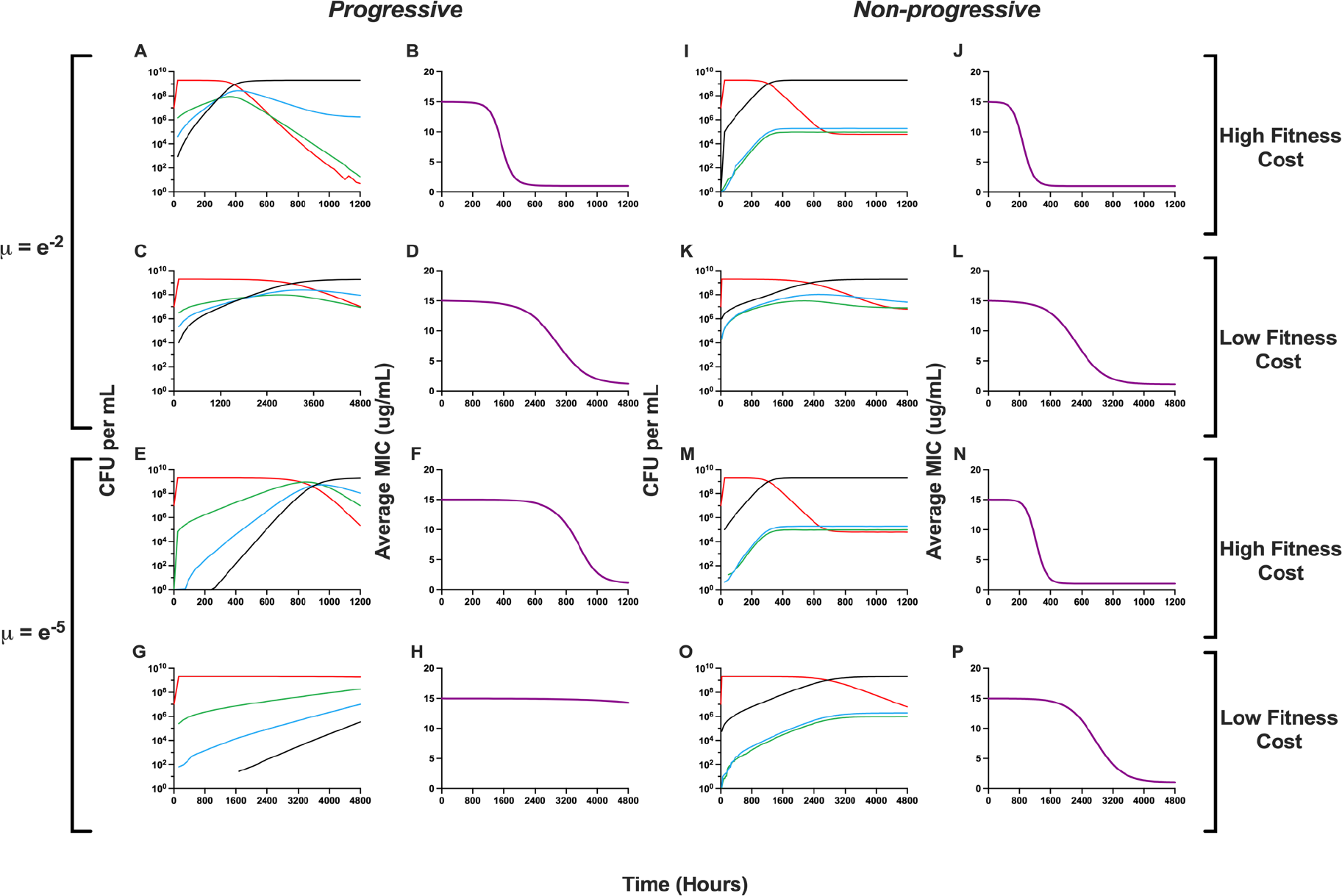
Response of the two heteroresistant models to the removal of antibiotics. Changes in the densities of the susceptible and resistant populations in the absence of the antibiotic and changes in the average MIC. S (black), R1 (blue), R2 (green), and R3 (red). Simulations with the high fitness cost were run for 1200 hours (50 days), while simulations with the low fitness cost were run for 4800 hours (200 days).

In the absence of antibiotics, the populations become increasingly dominated by more susceptible populations for both the progressive and non-progressive models of HR. This change in the composition of the populations is also reflected in a decline in the average MIC, approaching the level of the susceptible population. With the same fitness parameter and transition rates between states, µ, the rate of return to the susceptible state is greater for the non-progressive model than the progressive model. For both models, the rate of return to the sensitive state is proportional to the transition rate between states, µ, and the relative fitness cost of resistance. Notably, the new apparent equilibria obtained for both models differ substantially. In the progressive model, the most resistant populations are in continuous decline and will ultimately be lost or nearly so, while in the non-progressive model, all resistant populations are present at roughly equal frequency and appear to be in equilibrium. Of note is the vast difference in the time needed for the susceptible population to come to dominate; we list these times in Supplemental Table 2.

## Discussion

To elucidate the factors that govern the response of heteroresistant populations to antibiotics, we use mathematical and computer-simulation models to explore quantitatively: i) the factors responsible for generating the distribution of resistant sub-populations, ii) the response of heteroresistant populations to different concentrations of antibiotics, and iii) the amount of time required for an antibiotic-resistant heteroresistant population to become susceptible again when the treating antibiotic is removed.

We consider two models of heteroresistance (HR), which we call progressive and non-progressive. In both models, there are one or more sub-populations with different levels of resistance. In the progressive model, the more susceptible state transitions sequentially to the more resistant states, which in turn transition back to the less resistant states in the same sequence. In the non-progressive model, the susceptible population transitions directly to all the resistant states from the susceptible state before transitioning back directly to the susceptible state. In both models, the transition rates between states and the relative fitness cost of being resistant determine the distribution of the resistant populations in the absence of and in response to antibiotics.

The difference in the distribution of resistant states between these models with the parameters used is apparent with a Population Analysis Profile (PAP) test. With the progressive model, there are different resistance levels with distinct relative densities, which decline as the concentration of the antibiotic increases. With the non-progressive model, although there are multiple sub-populations with different levels of resistance, they likely would not appear as separate populations in a PAP test. The PAP test of the non-progressive HR looks more like that which would obtain with only two resistance levels, sensitive and resistant. However, the PAP tests are insufficient to differentiate the two models of HR, as there are conditions where non-progressive HR would look progressive (Supplemental Figure 7).

The models are also distinct in how they respond to antibiotics. In the progressive model, if the drug concentration is above the MIC of any of the sub-populations and the transition rate is low, the most resistant population can fail to emerge and ascend; this is true even though the drug concentration is still less than the MIC of the most resistant population. With the non-progressive model, the highest level of resistance will always emerge, no matter the transition rate. There are also differences in the population dynamics of each model when the antibiotics are removed. In the progressive model, the average MIC will return to that of the most susceptible population, and the most resistant populations will be lost. In contrast, in the non-progressive model, the average MIC will not decrease to that of the most susceptible population, and all the resistant sub-populations will remain present. One implication of this is when confronted with antibiotics, a heteroresistant population which is non-progressive will respond to the drugs more consistently and more rapidly than a progressive heteroresistant population.

The standard for detecting and defining a strain as heteroresistant is the PAP test which requires sub-populations to be more frequent than 10^-7^ and to have an MIC >8x that of the susceptible main population (9). These tests are cumbersome, costly, and are unlikely to be performed in clinical microbiology labs. Most critically, our results demonstrate that the PAP test is not sufficient to detect HR. There are conditions with the progressive and non-progressive models where populations would fail to meet the criteria set by the PAP test but would still survive confrontation with high doses of antibiotics – a false negative. There are also conditions where stable resistance would meet the HR criteria set by the PAP test – a false positive (Supplemental Figure 8). Moreover, there are conditions that would be called HR despite requiring thousands of hours to return to a sensitive MIC. To address this issue, although not in the current definition of HR, we recommend revisiting this definition to include the rate at which the MIC of potentially heteroresistant strains return to a susceptible MIC (12). This is especially important, as the epidemiological risk of HR is the rapid return to a seemingly sensitive state, and as demonstrated experimentally, this can happen in less than 50 generations for certain types of HR (12, 18, 19).

At this juncture, it is not clear how important HR is clinically even though animal experiments (20, 21) and some clinical studies suggest that it can increase the risk of persistent bacteremia, lead to longer hospital stays, and increase mortality (12). We argue that within a single infected individual, the distinction between the emergence of stable resistance and HR is manifest in the risk of treatment failure. With both mechanisms, antibiotics can select for the ascent of resistant sub-populations which will result in reduced treatment efficacy or even treatment failure, likely leading clinicians to change the treating drug in both cases. This risk of treatment failure is probabilistic in HR, as it is in stable resistance, due to other factors not considered here such as the host’s immune system, the compartmental heterogeneity of infection, and the local antibiotic concentrations. Due to the combination of these factors, treatment of a heteroresistant strain with a drug for which it is HR, will not necessarily lead to treatment failure. One distinction between HR and stable resistance is the rapid reversion of a heteroresistant population from a resistant to a susceptible state. This reversion has an additional clinical implication when considering infection transmission between individuals. Should an individual be infected with bacteria that are stably resistant to a drug, that resistance would appear on an assay such as the VITEK, and the drug for which they are resistant would not be used. If that individual is infected with heteroresistant bacteria, it would initially appear sensitive to a treating drug, but resistance could rapidly ascend. Then if that individual passes the infection on to another individual, due to the transient nature of HR, that infection would once again appear susceptible to the drug and once again the wrong drug would be chosen to treat the infection.

Although this study is purely theoretical, the parameters used in these models can be estimated experimentally with different species of bacteria and antibiotics of different classes. The hypotheses generated herein can be tested *in vitro* and, most importantly, can be rejected. There exists evidence supporting these two classes of HR, primarily in the form of PAP tests of known heteroresistant strains as exemplified by data shown in Supplemental Figure 9. A key objective for future experimental work is to determine how the actual mechanisms that can generate an unstable heteroresistant phenotype relate to these theoretical models. At present, we know of two main mechanisms that can generate HR: (i) alterations in copy number of resistance genes or their regulators by either tandem amplifications and/or alterations in plasmid copy number (12, 18, 22, 23), and (ii) regular point mutations that occur at a high frequency (18, 24-26). It is likely that mutational HR is best described by the non-progressive model where instability and reversion to susceptibility is driven by compensatory mutations that concomitantly reduce the fitness costs of the resistance mutations and lead to the loss of resistance (27-29). For gene amplification mechanisms it is less clear which theoretical model best describes their behavior since these mechanisms could have properties compatible with either model alone or a combination of the two, depending on the actual mechanism by which the amplifications are formed and lost. Further experimental work is needed to clarify these points. Finally, an unstable and transient resistant minority population could potentially also be generated by other types of mechanisms than those presently identified, including inducible resistances, epigenetic changes, and gene conversion events (30). In the supplemental text and Supplemental Table 3, we provide examples of HR mechanisms across several drug classes, predict which model of HR would most accurately pertain, and discuss the clinical implications.

There are, of course, caveats to consider with our models. Firstly, our models are not mechanistic and do not consider the genetic basis of progressive versus non-progressive HR and, as mentioned above, there are likely cases where certain mechanisms (e.g. gene amplification) could look either progressive, non-progressive, or somewhere in between depending on their specific mechanistic properties. Secondly, our models only include three resistant states, and these resistant states either do not transition between each other (non-progressive) or transition sequentially (progressive) – two extreme cases. Lastly, as with all pharmacodynamic studies, some elements have been neglected from these models, as mentioned previously, the host’s immune system and the compartmental heterogeneity of infection such as biofilms and abscesses, as well as variation in local antibiotic concentrations, all of which prohibit *in vitro* models and studies from making solid clinical predictions. All in all, a clear next step would be to test these predictions *in vitro* and then move to an *in vivo* model system. Crucially, we need to develop an understanding of how the definition of HR matches with the clinical implications, specifically considering the frequency and MIC cutoffs previously defined.

## Materials and Methods

### Numerical Solutions (Simulations)

For our numerical analysis of the coupled, ordered differential equations presented (Equations 2-11) we used Berkeley Madonna with the parameters presented in Table S1. Copies of the Berkeley Madonna programs used for these simulations are available at www.eclf.net.

### Bacteria

*Enterobacter cloacae* Mu208 is a carbapenem-resistant isolate collected by the Georgia Emerging Infections Program Multi-site Gram-negative Surveillance Initiative and described previously (21). *Burkholderia cepacia* complex isolate JC8 is a cystic fibrosis patient isolate collected by the Georgia Emerging Infections Program Multi-site Gram-negative Surveillance Initiative. *Escherichia coli* MG1655 was obtained from the Levin Lab’s bacterial collection.

### Rifampin Population Analysis Profile tests

Single colonies of *E. coli* MG1655 were inoculated into 10 mL lysogeny broth (BD, USA, Product #244610) and grown overnight at 37°C with shaking. Cultures were serially diluted in saline and all dilutions (10^0^ to 10^-7^) plated on LB agar plates (BD, USA, Product #244510) containing 0, 1, 2, 4, 8, and 16 times the MIC of rifampin (Thermo Fisher, USA, Product #J60836.03). Plates were grown at 37°C for 48 hours before the density of surviving colonies was estimated.

### *Burkholderia* and *Enterobacter* Population Analysis Profile tests

Single colonies of *B. cepacia* complex isolate JC8 and *E. cloacae* Mu208 were inoculated into 1.5 mL Mueller-Hinton broth (BD, USA, Product #275730) and cultures were grown overnight at 37°C with shaking. Cultures were serially diluted in phosphate-buffered saline and 10 µL of each dilution was spotted on Mueller-Hinton agar (BD, USA, Product #225250) plates containing 0, 0·125, 0·25, 0·5, 1, 2, and 4 times the breakpoint concentration of each antibiotic. Antibiotics used were ticarcillin disodium (BioVision, USA, Product #B1536) with clavulanate potassium salt (Cayman Chemical Company, USA, Procut #19456), amikacin sulfate (AstaTech, USA, Product # 40003), colistin sulfate salt (Sigma-Aldrich, USA, Product # C4461), and fosfomycin disodium salt (TCI America, USA, Product # F0889). For Mu208 on fosfomycin, broth and agar included 25 µg/mL glucose-6-phosphate (Sigma-Aldrich, USA, Product #10127647001). Plates were maintained at 37°C overnight for Mu208 and for 36-60 hours for JC8.

## Acknowledgments

We thank Dr. Danielle Steed for her discussion of the clinical implications of this manuscript. The authors would also like to thank Jason Chen, generally.

## Funding Sources

BRL, DAA, and DSW thank the U.S. National Institute of General Medical Sciences for their funding support via R35GM136407, the U.S. National Institute of Allergy and Infectious Diseases for their funding support via U19AI158080-02, and the Emory University Antibiotic Resistance Center. FB acknowledges the support of CIBERESP (CB06/02/0053) from the Carlos III Institute of Health of Spain. The funding sources had no role in the design of this study and will not have any role during its execution, analysis, interpretation of the data, or drafting of this report. The content is solely the responsibility of the authors and do not necessarily represent the official views of the National Institutes of Health nor those of the Carlos III Institute of Health of Spain.

## Data Availability

The Berkeley Madonna programs used for these simulations are available at ECLF.net. All data are presented in this article or its supplementary materials.

## Notes

### Competing Interest Statement

The authors have declared no competing interest.

### Summary of Updates

Updated abstract and discussion.

## References

1. Bengtsson B, Greko C. Antibiotic resistance--consequences for animal health, welfare, and food production. Ups J Med Sci. 2014;119(2):96–102.

2. Murray CJL, Ikuta KS, Sharara F, Swetschinski L, Robles Aguilar G, Gray A, et al. Global burden of bacterial antimicrobial resistance in 2019: a systematic analysis. The Lancet. 2022;399(10325):629–55.

3. MacGowan AP, Wise R. Establishing MIC breakpoints and the interpretation of in vitro susceptibility tests. J Antimicrob Chemother. 2001;48 Suppl 1:17–28.

4. Humphries R, Bobenchik AM, Hindler JA, Schuetz AN. Overview of Changes to the Clinical and Laboratory Standards Institute Performance Standards for Antimicrobial Susceptibility Testing, M100, 31st Edition. J Clin Microbiol. 2021;59(12):e0021321.

5. Bardelli M, Padovani M, Fiorentini S, Caruso A, Yamamura D, Gaskin M, et al. A side-by-side comparison of the performance and time-and-motion data of VITEK MS. Eur J Clin Microbiol Infect Dis. 2022;41(8):1115–25.

6. Comité de l’Antibiogramme de la Société Française de Microbiologie report 2003. Int J Antimicrob Agents. 2003;21(4):364–91.

7. Ericsson HM, Sherris JC. Antibiotic sensitivity testing. Report of an international collaborative study. Acta Pathol Microbiol Scand B Microbiol Immunol. 1971;217:Suppl 217:1+.

8. Brown DF, Brown L. Evaluation of the E test, a novel method of quantifying antimicrobial activity. J Antimicrob Chemother. 1991;27(2):185–90.

9. Falagas ME, Makris GC, Dimopoulos G, Matthaiou DK. Heteroresistance: a concern of increasing clinical significance? Clin Microbiol Infect. 2008;14(2):101–4.

10. Hiramatsu K, Aritaka N, Hanaki H, Kawasaki S, Hosoda Y, Hori S, et al. Dissemination in Japanese hospitals of strains of Staphylococcus aureus heterogeneously resistant to vancomycin. Lancet. 1997;350(9092):1670–3.

11. El-Halfawy OM, Valvano MA. Antimicrobial heteroresistance: an emerging field in need of clarity. Clinical microbiology reviews. 2015;28(1):191–207.

12. Andersson DI, Nicoloff H, Hjort K. Mechanisms and clinical relevance of bacterial heteroresistance. Nature Reviews Microbiology. 2019;17(8):479–96.

13. Regoes RR, Wiuff C, Zappala RM, Garner KN, Baquero F, Levin BR. Pharmacodynamic functions: a multiparameter approach to the design of antibiotic treatment regimens. Antimicrobial agents and chemotherapy. 2004;48(10):3670–6.

14. Berryhill BA, Gil-Gil T, Manuel JA, Smith AP, Margollis E, Baquero F, et al. What’s the Matter with MICs: Bacterial Nutrition, Limiting Resources, and Antibiotic Pharmacodynamics. Microbiology Spectrum. 2023:e04091–22.

15. Monod J. The growth of bacterial cultures. Annual review of microbiology. 1949;3(1):371–94.

16. Metropolis N, Ulam S. The Monte Carlo Method. Journal of the American Statistical Association. 1949;44(247):335–41.

17. Wanner G, Hairer E. Solving ordinary differential equations II: Springer Berlin Heidelberg New York; 1996.

18. Nicoloff H, Hjort K, Levin BR, Andersson DI. The high prevalence of antibiotic heteroresistance in pathogenic bacteria is mainly caused by gene amplification. Nature Microbiology. 2019;4(3):504–14.

19. Pereira C, Larsson J, Hjort K, Elf J, Andersson DI. The highly dynamic nature of bacterial heteroresistance impairs its clinical detection. Communications Biology. 2021;4(1):521.

20. Band VI, Satola SW, Burd EM, Farley MM, Jacob JT, Weiss DS. Carbapenem-Resistant Klebsiella pneumoniae Exhibiting Clinically Undetected Colistin Heteroresistance Leads to Treatment Failure in a Murine Model of Infection. mBio. 2018;9(2).

21. Band VI, Hufnagel DA, Jaggavarapu S, Sherman EX, Wozniak JE, Satola SW, et al. Antibiotic combinations that exploit heteroresistance to multiple drugs effectively control infection. Nat Microbiol. 2019;4(10):1627–35.

22. Hjort K, Nicoloff H, Andersson DI. Unstable tandem gene amplification generates heteroresistance (variation in resistance within a population) to colistin in Salmonella enterica. Mol Microbiol. 2016;102(2):274–89.

23. Anderson SE, Sherman EX, Weiss DS, Rather PN. Aminoglycoside Heteroresistance in Acinetobacter baumannii AB5075. mSphere. 2018;3(4).

24. Charretier Y, Diene SM, Baud D, Chatellier S, Santiago-Allexant E, van Belkum A, et al. Colistin Heteroresistance and Involvement of the PmrAB Regulatory System in Acinetobacter baumannii. Antimicrob Agents Chemother. 2018;62(9).

25. Halaby T, Kucukkose E, Janssen AB, Rogers MR, Doorduijn DJ, van der Zanden AG, et al. Genomic Characterization of Colistin Heteroresistance in Klebsiella pneumoniae during a Nosocomial Outbreak. Antimicrob Agents Chemother. 2016;60(11):6837–43.

26. Jayol A, Nordmann P, Brink A, Poirel L. Heteroresistance to colistin in Klebsiella pneumoniae associated with alterations in the PhoPQ regulatory system. Antimicrob Agents Chemother. 2015;59(5):2780–4.

27. Andersson DI, Levin BR. The biological cost of antibiotic resistance. Curr Opin Microbiol. 1999;2(5):489–93.

28. Andersson DI, Hughes D. Antibiotic resistance and its cost: is it possible to reverse resistance? Nat Rev Microbiol. 2010;8(4):260–71.

29. Allen RC, Engelstädter J, Bonhoeffer S, McDonald BA, Hall AR. Reversing resistance: different routes and common themes across pathogens. Proc Biol Sci. 2017;284(1863).

30. Meka VG, Gold HS, Cooke A, Venkataraman L, Eliopoulos GM, Moellering RC, Jr., et al. Reversion to susceptibility in a linezolid-resistant clinical isolate of Staphylococcus aureus. J Antimicrob Chemother. 2004;54(4):818–20.

