## Supplemental materials for "Theoretical Considerations and Empirical Predictions of the Pharmaco- and Population Dynamics of Heteroresistance"

**Supplemental Materials for**
**Theoretical considerations and empirical predictions about the pharmaco- and population dynamics of antibiotic heteroresistance**

**This 12 page PDF file includes:**

Supplemental Text
Figures S1 to S9
Tables S1 to S3

**Supplemental Text**

By definition, a heteroresistant strain of bacteria contains one or more resistant sub-populations at a high frequency (greater than $10^{-7}$) and with an MIC at least 8x higher than the dominant, susceptible population. For non-progressive HR, all resistant subpopulations emerge by transition directly from the susceptible population. In the progressive model of HR, several sequential transitional steps are required to generate the most resistant population. These resistant subpopulations grow in the presence of the antibiotic, making this phenomenon distinct from persistence. Therefore, HR could have a greater clinical impact on antibiotic treatment than persistence. In most cases, HR remains under-detected by routine susceptibility testing. In Supplemental Table 2 we present examples of genes and mechanisms by which HR may emerge for eleven different classes of antibiotics. It should be noted that the response of one bacteria to one antibiotic could fit both the progressive and non-progressive HR models (Supplemental Figures 5 and 7).

As predicted by our simulated PAP tests and observed in previous data, the frequency of some heteroresistant sub-populations are below the $10^{-7}$ frequency. Though these strains may be below the definitional threshold, they are still of clinical interest, since the absolute number of cells in some infection sites can easily surpass $10^{7}$ cells.

In principle, the transition rate both to and from resistance in both models would depend on the mechanism, underlying fitness cost of being resistant, and frequency of fitness cost compensatory mutations. In Supplemental Table 2, we consider the potential rates both to and from resistance based on mechanism and fitness cost.

The difference in mechanism and transition rates leads us to propose different antibiotic treatment regimens to deal with either progressive or non-progressive heteroresistant infections. For an infection with a non-progressive heteroresistant bacteria, wherein a resistant sub-population could rapidly replace the sensitive population, we suggest frequent (optimally, daily) surveillance of the susceptibility of the bacteria. After a successful therapy, in the case of relapse, susceptibility may be rapidly restored and the same antibiotic could be reconsidered for therapy. We, of course, recommend ensuring that susceptibility has been restored before reusing the same antibiotic. For an infection with a progressive heteroresistant bacteria, it is critical to prevent the evolution of the first resistant state. This may be done by increasing the dose of the antibiotic. In the case of progressive HR, susceptibility testing need not be done as frequently as with the non-progressive bacteria. Should treatment fail and the infection relapse, the same antibiotic should not be used as the transition from resistant to sensitive is expected to take a longer time than in the case of a non-progressive heteroresistant infection. In general, if an infecting bacteria is shown to be heteroresistant, antibiotic combinations should always be considered, especially in high-risk patients.

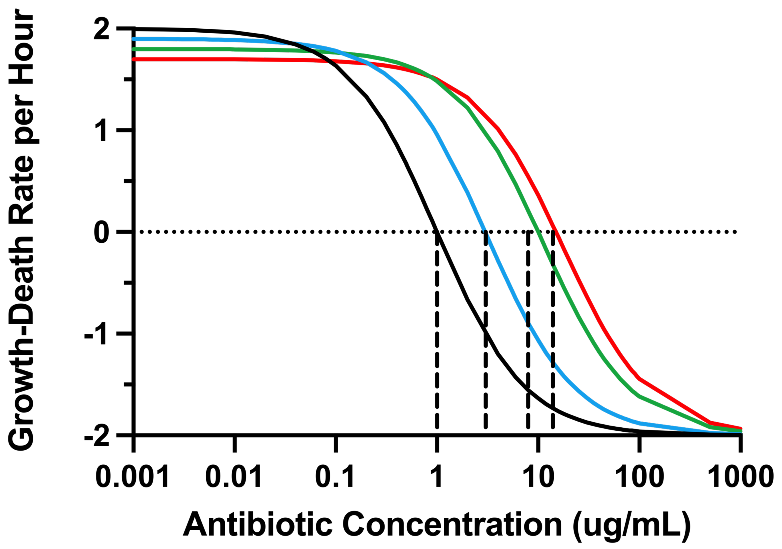


**Figure S1. Hill function pharmacodynamics**. The relationship between the antibiotic concentration and the rate of growth and death of the bacteria. The four bacterial populations, S (black), R1 (blue), R2 (green), and R3 (red) differ in their maximum growth rates, $v_S$ =2·0, $v_1$ =1·9. $v_2$=1·8, $v_3$=1·7 per cell per hour, respectively. These bacteria also differ in the resistance levels as measured by their respective MICs, $MIC_S$ =1·0, $MIC_{R1}$=3·0, $MIC_{R2}$=8·0, $MIC_{R3}$=15·0 μg/mL, respectively. All the bacteria have the same negative growth rate, $v_{MIN}$ =-2·0 per cell per hour and the same Hill coefficient, κ=1·0. For the depicted pharmacodynamics, we assume that there is no limitation in the concentration of the resource.

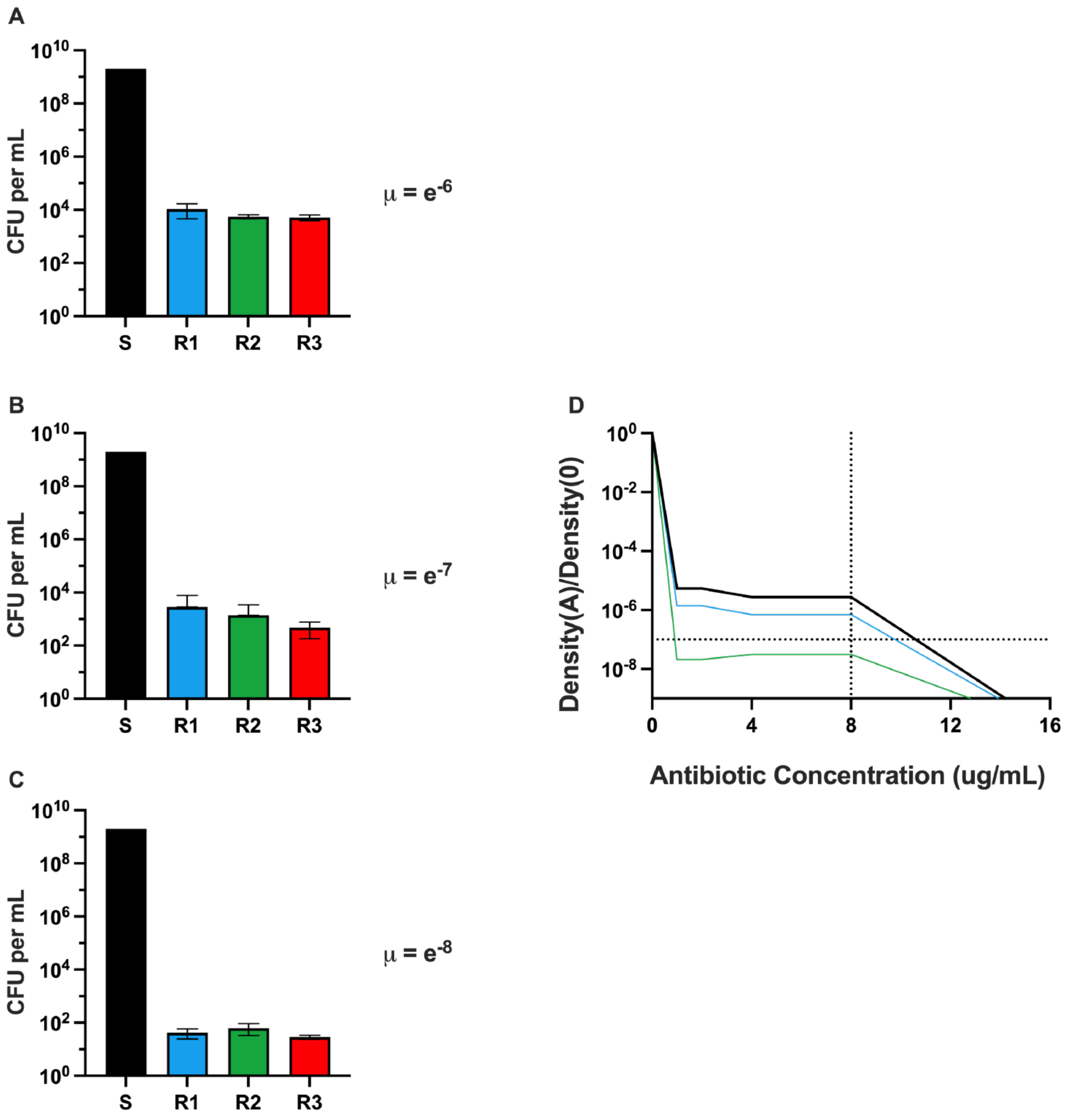


**Figure S2. Non-Progressive model with low transition rates.** (A-C) The distribution of stationary phase densities when grown up from a single cell of S with three different transition rates. A: $\mu=10^{-6}$, B: $\mu=10^{-7}$, C: $\mu=10^{-8}$. (D) A corresponding PAP test when $\mu=10^{-6}$ (Black), $\mu=10^{-7}$ (Blue), and $\mu=10^{-8}$ (Green). Parameters used in this figure: maximum growth rates, $v_S=2{\cdot}0$, $v_{R1}=1{\cdot}9$, $v_{R2}=1{\cdot}8$, $v_{R3}=1{\cdot}7$ per cell per hour, $MIC_S=1{\cdot}0$, $MIC_{R1}=3{\cdot}0$, $MIC_{R2}=10{\cdot}0$, and $MIC_{R3}=15$ μg/mL. S (black), R1(blue), R2 (green), R3 (red).

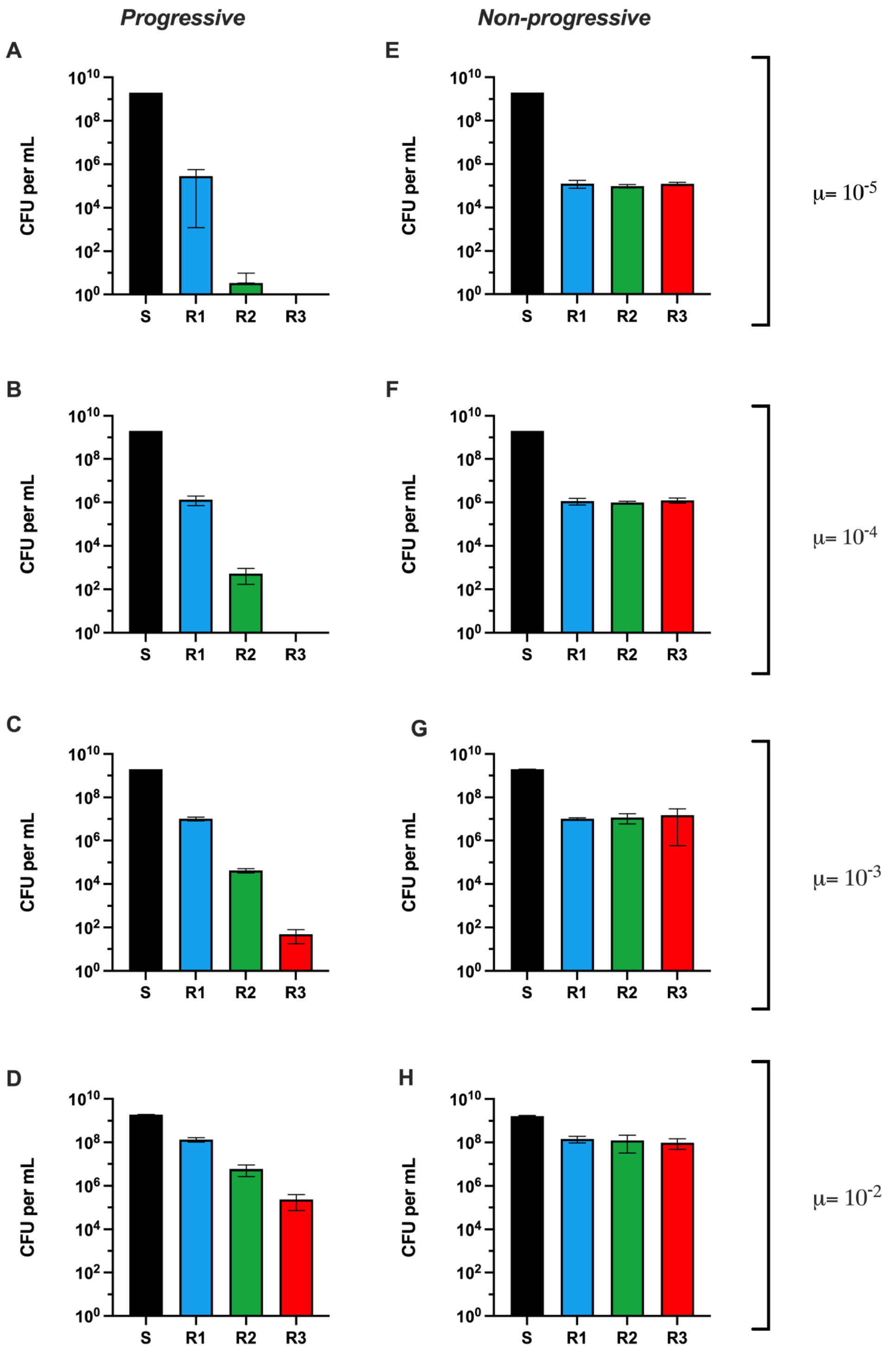


**Figure S3. Distribution of stationary phase densities with grown up from a single S cell with a low fitness cost**. Shown on the left (A, B, C, and D) are the means and standard deviations of the stationary phase densities of the S (black), R1 (blue), R2 (green), and R3 (red) subpopulations from five independent runs with the progressive model with different transition rates, μ=$10^{-5}$, $10^{-4}$, $10^{-3}$, and $10^{-2}$ per cell per hour for A, B, C, and D, respectively. On the right (E, F, G, and H) are the corresponding distributions for runs made with the non-progressive model with these respective transition rates. Unlike in the main body, this figure assumes a low fitness cost with maximum growth rates, $v_S$=2·0, $v_{R1}$=1·99, $v_{R2}$=1·98, and $v_{R3}$=1·97 per cell per hour.

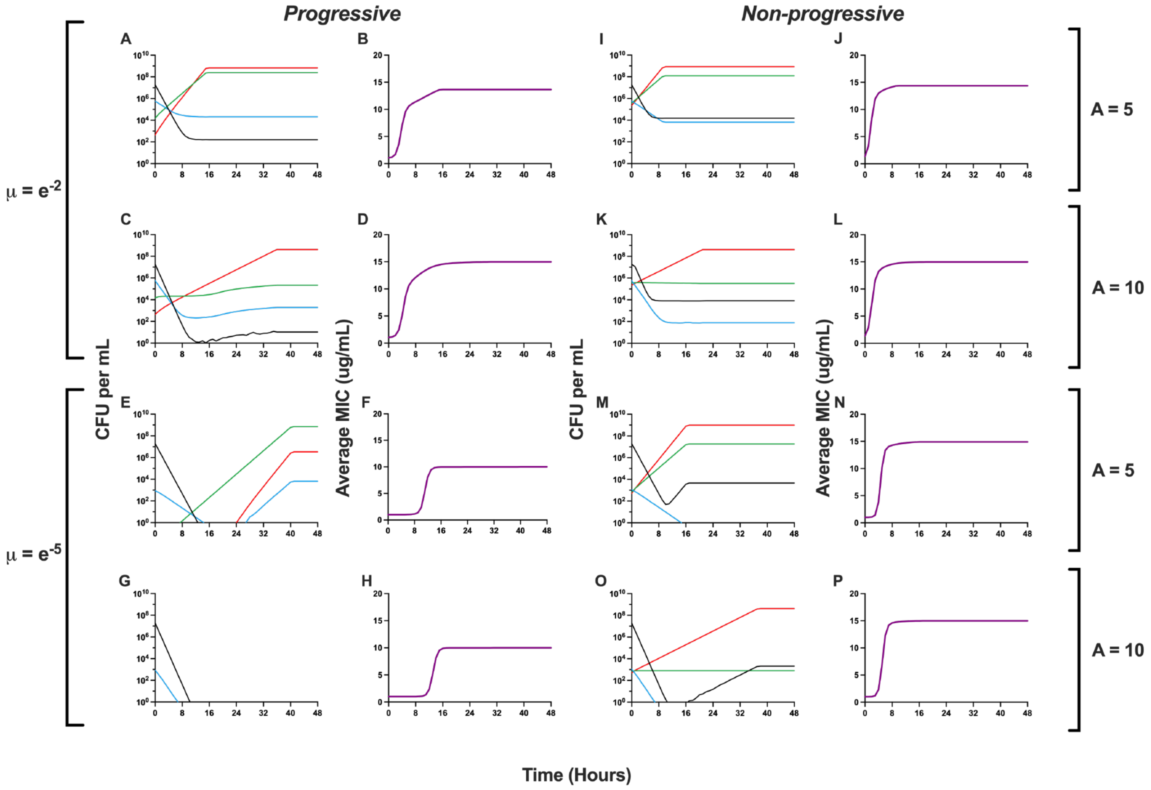


**Figure S4. Response of heteroresistant populations to two concentrations of antibiotics.** Changes in the densities of the susceptible and the different resistant populations and the average MIC where μ=$10^{-2}$ or μ=$10^{-5}$ per cell per hour when exposed to either 5 μg/mL or 10 μg/mL (5x and 10x the MIC of the susceptible main population) of the antibiotic. S (black), R1 (blue), R2 (green), and R3 (red). The initial densities for the progressive and non-progressive simulations are 1/100 of the densities presented in Figure 3D and A and Figure 3H and E respectively with the given transition rate.

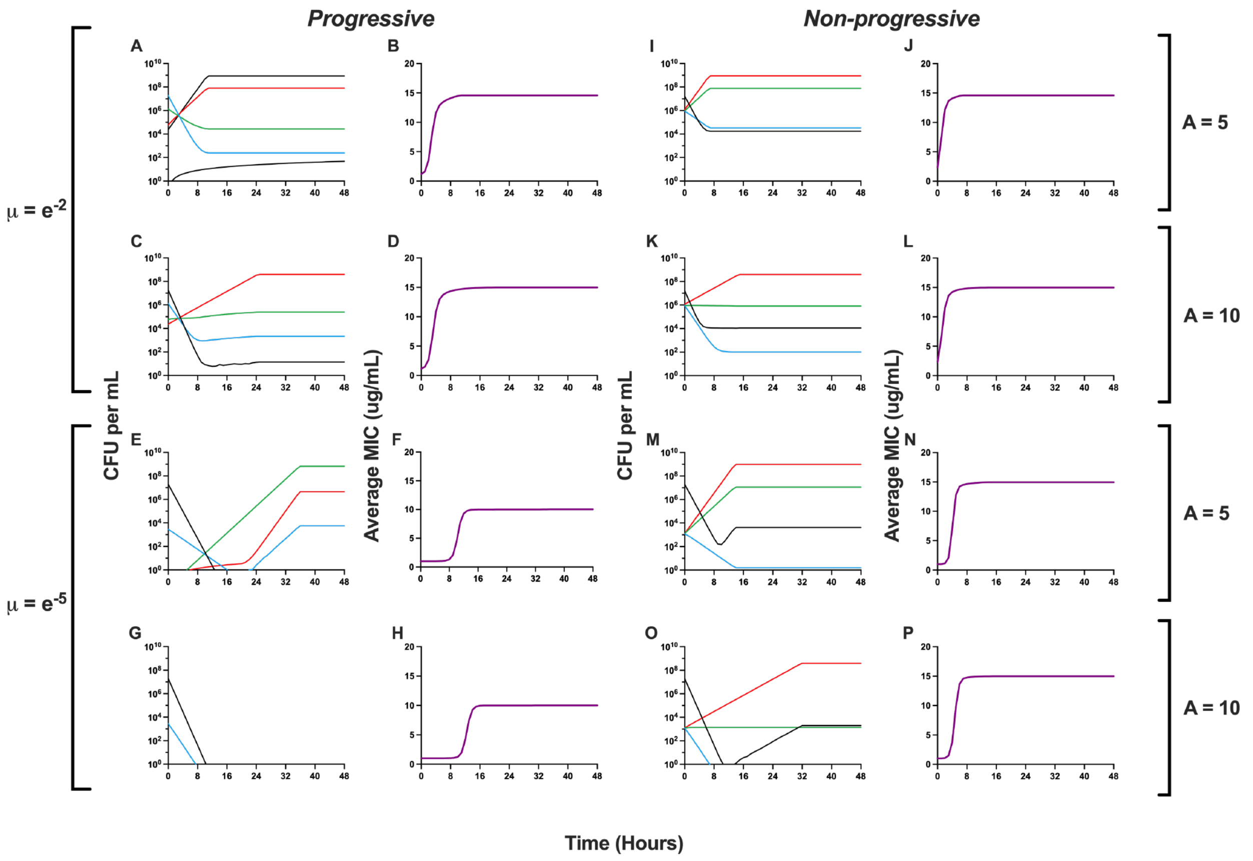


**Figure S5**. **Response of heteroresistant populations to different concentrations of antibiotics with a low fitness cost.** Changes in the densities of the sensitive and different resistant populations and the average MIC where $\mu=10^{-2}$ or $\mu=10^{-5}$ per cell per hour when exposed to either 5 μg/mL or 10 μg/mL of the antibiotic. S (black), R1 (blue), R2 (green), and R3 (red). Low fitness cost maximum growth rates, $v_S=2{\cdot}0$, $v_{R1}=1{\cdot}99$, $v_{R2}=1{\cdot}98$, and $v_{R3}=1{\cdot}97$ per cell per hour**.**

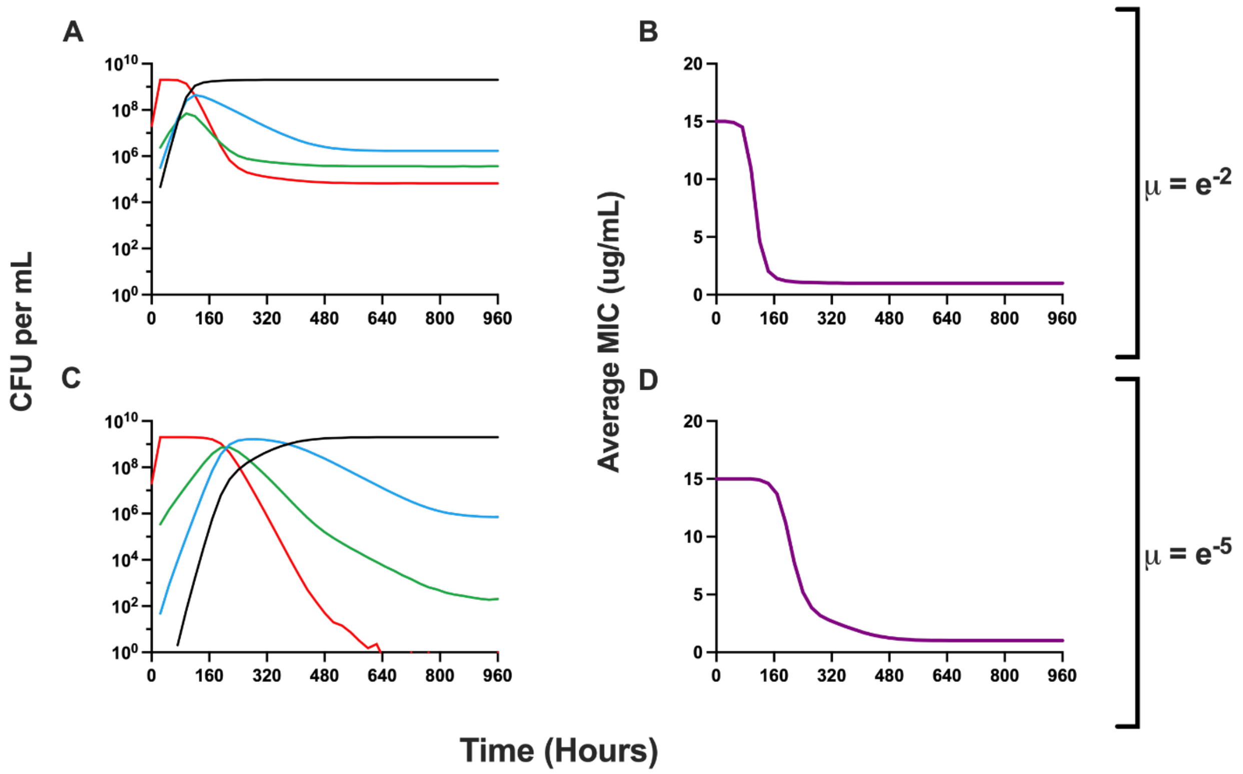


**Figure S6**. **Response of the two heteroresistant models to the removal of antibiotics with a very high fitness cost.** Changes in the densities of the susceptible and resistant populations in the absence of the antibiotic and changes in the average MIC. S (black), R1 (blue), R2 (green), and R3 (red). Simulations with the very high fitness cost were run for 500 hours (approximately 20 days). The return to sensitive when $\mu=10^{-2}$ occurs at 110 hours (approximately 5 days) and when $\mu=10^{-5}$ it takes 370 hours (approximately 15 days).

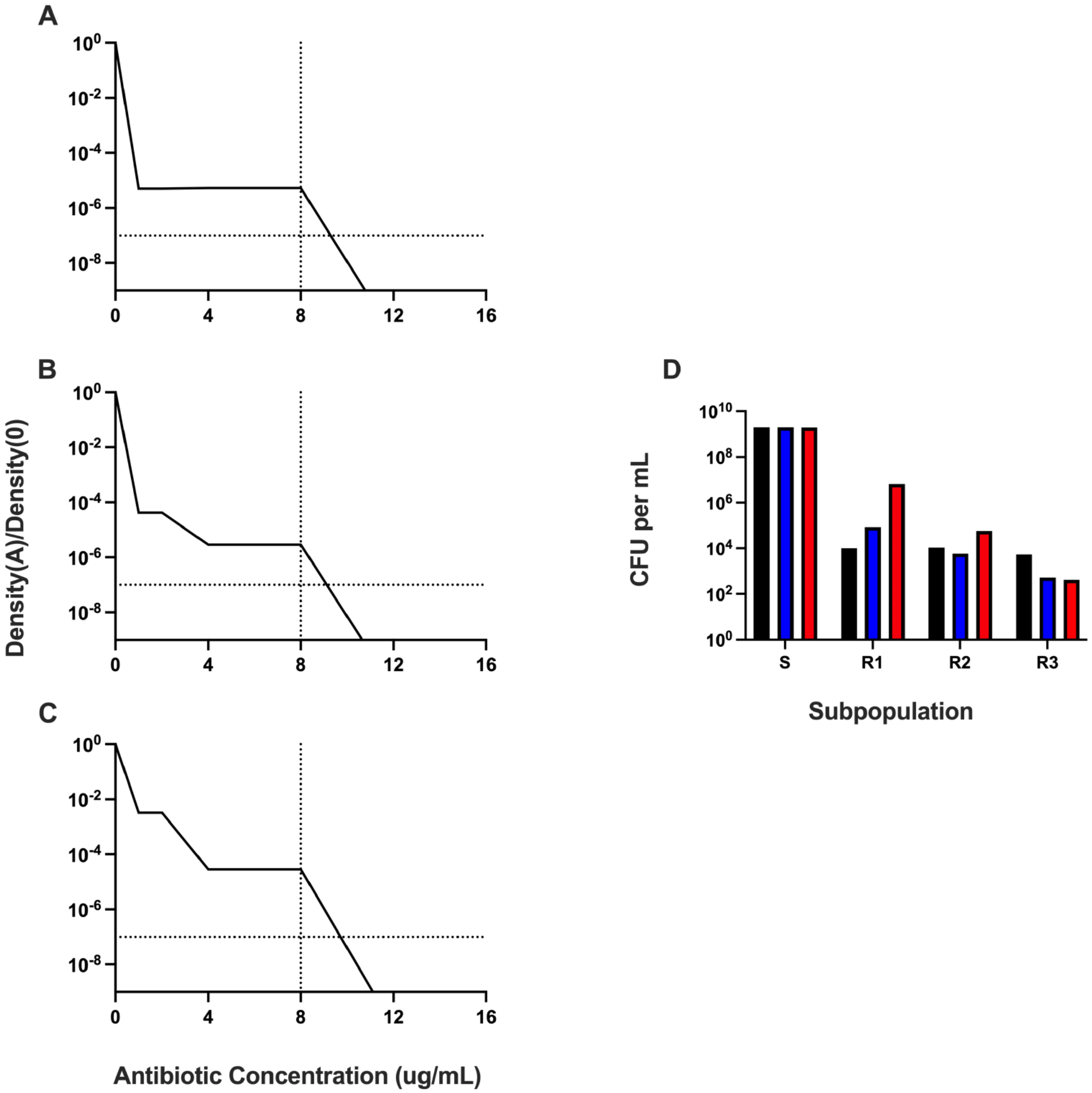


**Figure S7. Population analysis profiles for the non-progressive model with different transition rates between states.** (A) PAP test with transition rates $\mu=10^{-6}$ per cell per hour. (B) PAP test with transition rates $\mu_{SR1}=\mu_{R1S}=10^{-5}$, $\mu_{SR2}=\mu_{R2S}=10^{-6}$, and $\mu_{SR3}=\mu_{R3S}=10^{-7}$ per cell per hour. (C) PAP test with transition rates $\mu_{SR1}=\mu_{R1S}=10^{-3}$, $\mu_{SR2}=\mu_{R2S}=10^{-5}$, and $\mu_{SR3}=\mu_{R3S}=10^{-7}$ per cell per hour. (D) Distribution of stationary phase densities when grown up from a single cell of S with the transition rates from panel A (black), panel B (blue), and panel C (red). Parameters used in these simulations: maximum growth rates, $v_S=2{\cdot}0$, $v_{R1}=1{\cdot}9$, $v_{R2}=1{\cdot}8$, $v_{R3}=1{\cdot}7$ per cell per hour, $MIC_S=1{\cdot}0$, $MIC_{R1}=3{\cdot}0$, $MIC_{R2}=10{\cdot}0$, and $MIC_{R3}=15$ μg/mL.

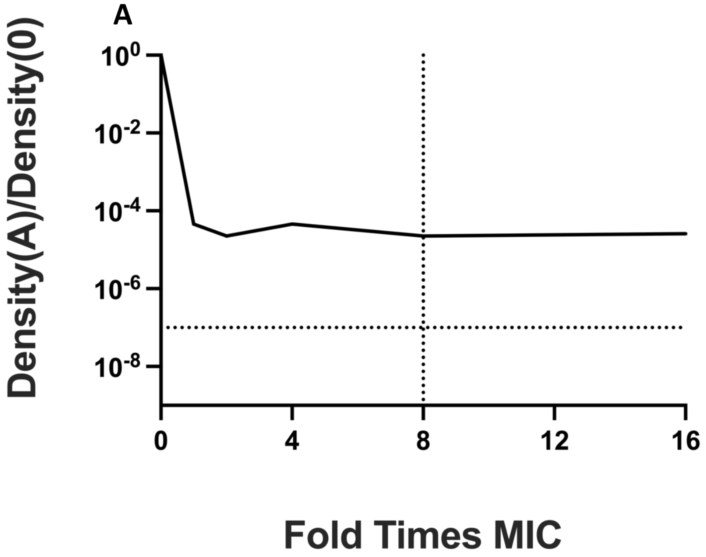


**Figure S8. Population Analysis Profiles of *Escherichia coli* with rifampin result in false positives.** *E. coli* MG1655 stable resistant mutants appearing heteroresistant to rifampin.

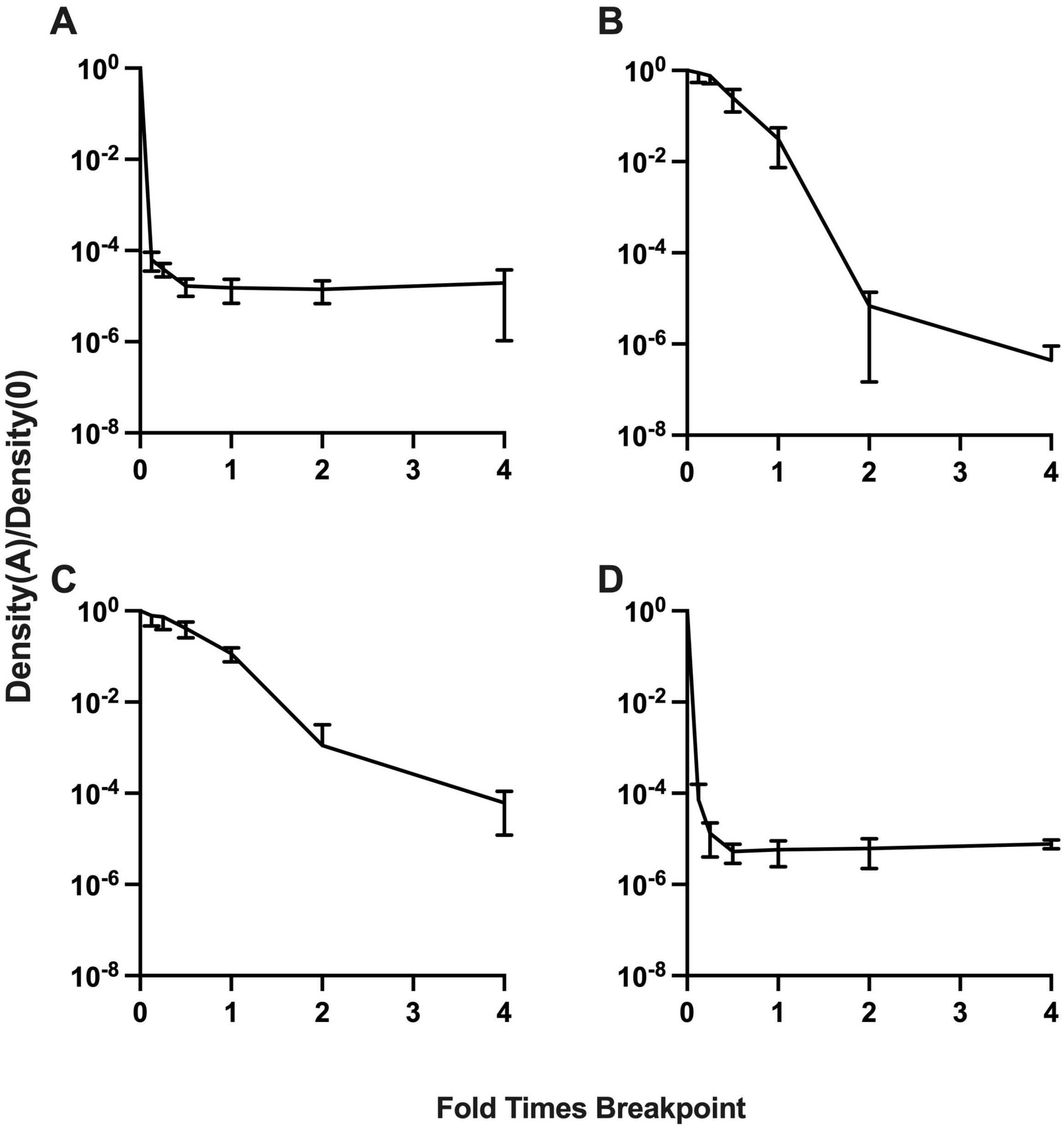


**Figure S9. Emperical population analysis profiles.** (A) *Burkholderia cepacia* JC8 complex heteroresistant to ticarcillin/clavulanate. (B) *B. cepacian* JC8 complex heteroresistant to amikacin. (C) *Enterobacter cloacae* Mu208 heteroreistant to fosfomycin. (D) *E. cloacae* Mu208 heteroresistant to colistin. Shown are means and standard deviations for each PAP test. Breakpoints used were: 128/2 μg/mL ticarcillin/clavulanate, 64 μg/mL amikacin, 256 μg/mL fosfomycin, and 4 μg/mL colistin.

**Supplemental Table 1. Parameters used in the simulations**

| Parameter | Definition and Dimensions | Values |
|---|---|---|
| $v_s$, $v_{R1}$, $v_{R2}$, $v_{R3}$ | Maximum growth rates Very high<br>(per cell/hr) High<br>Low | 2.0, 1.7, 1.5, 1.2<br>2.0, 1.9, 1.8, 1.7<br>2.0, 1.99, 1.98, 1.97 |
| $\mu_x$ | Transition rates (per cell/hr) | $10^{-2}$,$10^{-3}$, …, $10^{-8}$ |
| $v_{MIN}$ | Minimum growth/death rates (per cell/hr) | -2.0 |
| k | Hill coefficient | 1.0 |
| $MIC_S$, $MIC_{R1}$, $MIC_{R2}$, $MIC_{R3}$ | MIC (µg/mL) | 1.0, 3.0, 10.0, 15.0 |
| k | Monod constant | 1.0 |
| e | Resource conversion efficiency (µg) | 5 x 10-$^{7}$ |
| C | Maximum resource (µg/mL) | 1000 |
| A | Antibiotic concentration (µg/mL) | 0, 5, 10 |
| r | Resource concentration at a given time (µg) | $0 \leq r \leq 1000$ |

**Supplemental Table 2. Time for the heteroresistant populations to return to sensitivity**

| Condition (Panel of Figure 6) | Approximate time in hours (days) before the sensitive population dominates |
|---|---|
| Progressive Model | |
| A | 400 (16.7 days) |
| C | 3000 (125 days) |
| E | 900 (37.5 days) |
| G | 8000 (333 days) |
| Non-progressive Model | |
| I | 250 (10.4 days) |
| K | 2400 (100 days) |
| M | 300 (12.5 days) |
| O | 2800 (117 days) |

**Supplemental Table 3. Examples* of non-progressive and progressive heteroresistance in different antibiotic classes**

| Antibiotic Class | Non-progressive HR | Progressive HR | Transition to Resistance | Transition to Susceptbility |
|---|---|---|---|---|
| **Quinolones** | • *gyrA* mutations (nalidixic acid) | • *gyrA* mutations (fluoroquinolones)<br>• *gyrA+parC* mutations (fluoroquinolones)<br>• *gyrA* and efflux pumps amplifications (nalidixic acid) | Fast | Slow, but fast in amplification |
| **Aminoglycosides** | • *rpsL* mutations (streptomycin)<br>• *16S rRNA* A1402G mutation (kanamycin, gentamicin, and spectinomycin) | • *aadB* amplifications<br>• *aac*(3)**II**d amplifications | Fast | Slow, but fast when either RpsL K43T compensory mutation occurs for streptomycin or in amplification events |
| **Beta-lactams** | • Penicillin binding proteins (PBPs)<br>• Outer membrane porin o*prD* (carbapenems) | • Beta-lactamases *bla*TEM-1, TEM-10, and TEM-12 mutations<br>• Beta-lactamase amplication<br>• PBPs mutations<br>• *oprD* mutations in combination with efflux pump mutations (carbapenems) | Fast | Fast or slow, depending on associated fitness cost |
| **Macrolides** | • *23S rRNA* mutations<br>• L4, L22 ribosomal mutations | • Overproduction or mutations in efflux pumps<br>• Progressive predominance of L4, L22 mutated ribosomes | Fast with low rRNA copy number | Fast due to high fitness cost |
| **Rifampin** | • *rpoB* mutations | | Fast | Fast due to *tufA* or *rpoB* Y526H compensatory mutations |
| **Trimethoprim** | • *folA* mutations | • Stepwise *folA* mutations | Fast | Slow |
| **Linezolid** | | • Single nucleotide transversion in *23S rRNA* (typically G2576) leading to recombination with other copies of the gene (gene conversion) | Slow | Fast |
| **Fusidic acid** | • *fusA* mutations | | Fast | Fast |

| | | | | |
|---|---|---|---|---|
| **Lipo and glycopeptides** | | • Mutations in *mprF*, *yycFG* operon and genes encoding subunits of RNA polymerase<br>• Amplification of *pmrD*, regulating genes encoding proteins that modify lipid A | Slow | Fast due to high fitness cost |
| **Mupirocin** | • *ileS* mutations | • Consecutive mutations in *ileS* | Fast | Slow |
| **Fosfomycin** | • Mutations in *murA, uhpT,* and *glpT* | | Fast (in the absence of glucose-6 phosphate) | Fast |
| **Tigecycline Tetracyclines** | | • Overproduction of efflux pumps AcrAB–TolC, OqxAB, and MacAB<br>• Sequential mutations in *ramR*, *lon*, and *rpsJ*<br>• Amplification of *tet(A)* | Fast | Fast |

* Details and further references about above examples of mentioned resistance mechanisms can be found in: Baquero F, Martínez JL, F Lanza V, Rodríguez-Beltrán J, Galán JC, San Millán A, Cantón R, Coque TM. (2021) Evolutionary Pathways and Trajectories in Antibiotic Resistance. *Rev Clin Microbiol*. 34(4):e0005019, and Pereira, C., Larsson, J., Hjort, K., Elf, J., Andersson, D. I. (2021). The highly dynamic nature of bacterial heteroresistance impairs its clinical detection. *Communications Biology*, *4*(1), 521.
